# KiMA: Kinematic Motion Analysis for Spinal Cord Injury Research

**DOI:** 10.64898/2026.08.05.742960

**Authors:** Manojkumar Kumaran, N. R. Siva Shanmugam, Ishwariya Venkatesh

## Abstract

Accurate quantification of locomotor recovery is essential for evaluating therapeutic outcomes in spinal cord injury (SCI) models. Manual scoring systems remain observer-dependent, and commercial gait-analysis platforms are costly and proprietary. Markerless pose-estimation tools such as DeepLabCut generate accurate body-part coordinates, but converting these coordinates into biologically meaningful locomotor parameters typically requires custom programming and multiple external tools. We developed KiMA (Kinematic Motion Analysis), an open-source, browser-based suite for integrated analysis of rodent gait and hindlimb kinematics. KiMA accepts DeepLabCut coordinate files and performs automated coordinate parsing, stick-figure reconstruction, frame-by-frame movement inspection, and single- and multi-sample analysis, with dedicated workflows for ladder and rung analysis, footfall detection, and CatWalk gait analysis. The platform quantifies joint angles (metatarsophalangeal, ankle, knee, hip, and pelvic), stride length, stride width, cadence, stance and swing durations, paw-contact events, swing clearance, and locomotor symmetry, and supports cohort-level comparisons, correlation analysis, principal component analysis, and export of processed datasets and publication-quality figures. Because KiMA runs entirely within a standard web browser, it requires no software installation or local programming environment, supporting cross-platform accessibility and data privacy. By unifying gait quantification, visualization, and multivariate analysis in a single interface, KiMA lowers the computational barrier to markerless locomotor analysis and helps researchers detect subtle functional recovery after SCI.

## Introduction

Quantifying locomotor behavior remains a bottleneck in translational spinal cord injury (SCI) research. In murine models, spinal cord damage from contusion, transection, or compression at thoracic or lumbar levels disrupts descending motor tracts and produces varying degrees of hindlimb paralysis and motor coordination deficits [1]. Because the goal of regenerative therapies is restoration of motor function, objective quantification of hindlimb locomotion is essential for evaluating spinal cord repair, motor neuron survival, and axon regeneration [2].

Quantitative gait analysis in SCI research has historically relied on manual scoring systems, most notably the Basso Mouse Scale (BMS) [1] and the Basso-Beattie-Bresnahan (BBB) rating scale [2]. These scales provide useful qualitative readouts of joint movement, weight support, and coordination, but they are labor-intensive, susceptible to observer bias, and insensitive to subtle kinematic changes. Commercial automated systems such as CatWalk XT [3] and DigiGait [4] offer objective alternatives, yet their high acquisition costs and proprietary software limit widespread adoption and constrain the detection of fine-grained behavioral improvements in SCI.[5]

Markerless pose-estimation frameworks such as DeepLabCut [6] and SLEAP [7] have expanded what is tractable in behavioral neuroscience, enabling automated tracking of anatomical landmarks (iliac crest, hip, knee, ankle, metatarsophalangeal joint, toe) with pixel-level precision. These systems, however, output raw coordinates rather than interpretable locomotor parameters. Converting raw coordinates into standard kinematic measures such as joint angles, stance and swing durations, stride lengths, and cadence requires custom code, which is a barrier for researchers without extensive programming experience.

Downstream toolboxes such as ALMA (Automated Limb Motion Analysis) [8] address part of this gap, but effective use of these local Python packages still requires computational training that many experimental biologists do not have. Most existing open-source toolboxes also lack integrated statistical testing, so users must export processed data to third-party software (GraphPad Prism, R, Python) for group comparisons. The result is a fragmented pipeline in which visualization, quantification, and statistics live in separate environments.

To address these limitations, we developed KiMA (Kinematics Motion Analysis), a zero-installation web platform for evaluating motor recovery in SCI models. KiMA runs entirely in a standard web browser, allowing researchers to import DeepLabCut coordinate files and extract kinematic parameters without local software setup or programming expertise. It provides dedicated modules for the Horizontal Ladder & Rung test, Foot Fall analysis, CatWalk-style overground gait, and Treadmill locomotion. KiMA also includes a built-in statistical engine (Student’s t-test, one-way ANOVA), correlation matrix visualization, and multivariate analysis (Principal Component Analysis, Random Forest classification), consolidating quantification, visualization, and inference into a single pipeline for characterizing functional recovery after SCI.

## Methods

### KiMA toolbox

We developed KiMA as a web-based, zero-installation toolbox for multi-modal analysis of gait, footfall, and joint kinematics in mouse models of spinal cord injury. The toolbox provides a graphical user interface (GUI) with four core functionalities: (i) automated joint-coordinate parsing and stick-figure reconstruction from DeepLabCut output; (ii) task-specific analysis modules for Ladder & Rung, Foot Fall, CatWalk, and Treadmill locomotion; (iii) built-in cohort statistics and dimensionality reduction (Student’s t-test, one-way ANOVA, Pearson correlation, and Principal Component Analysis); and (iv) high-resolution export of gait kinematics and summary figures. KiMA is an open-source, client-side web application built with HTML5 and JavaScript, so all computation runs locally in the browser without a Python environment or server backend. This design supports cross-platform use and keeps raw data on the user’s machine.

Parameters extracted by KiMA include dynamic joint angles (toe, MTP, ankle, knee, hip, and iliac crest), footprint contact events, stance and swing durations, stride length and stride width, cadence, swing clearance height, and unilateral Symmetry Indices (SI) for locomotor side bias. After parameter extraction, the built-in statistics module computes group comparisons, Pearson correlation heatmaps to identify covarying features, and PCA projections for visualizing group separation. The GUI additionally provides a timeline view for frame-by-frame auditing of tracked joint movements, which is useful for validating fine motor tasks such as ladder-rung traversal. Full documentation, the live application, and source code are available at https://kimasuite.in and https://github.com/mano2991/KiMA.

### Video acquisition and DeepLabCut model training

Behavioral videos were acquired with an iphone 12 Pro camera at 1080p and 30 fps, using a fixed focal length of [26 mm] and a camera-to-apparatus distance of 45 cm, positioned parallel to the apparatus. Approximately 500 frames per trial were used for network training and evaluation, drawn from across experimental stages (uninjured, 1 WPI, 3 WPI) and animals to capture pose variability. Frame-by-frame markerless pose estimation was performed using DeepLabCut v3 [6].

For the Ladder & Rung module, six hindlimb landmarks were manually labelled on each training frame: iliac crest, hip, knee, ankle, metatarsophalangeal (MTP) joint, and toe. For the Foot Fall and CatWalk modules, four limb endpoints were labelled: left forelimb (LF), right forelimb (RF), left hindlimb (LH), and right hindlimb (RH). A total of 50 representative frames were extracted and manually labelled per video, with a 70:30 train-to-test split applied within DeepLabCut.

Networks were built on a ResNet-50 backbone initialised with ImageNet pre-trained weights. Each module-specific network was trained for 1,000,000 iterations until the cross-entropy loss plateaued and per-landmark test error stabilised below 2.8 pixels (mean test RMSE, corresponding to < 1.0 mm physical spatial accuracy). Training and inference were performed on a workstation with an Intel Xeon 64-core processor, 512 GB RAM, and an NVIDIA GeForce RTX 2080 Ti GPU (12 GB VRAM).

### Random Forest classification

To test whether the multivariate kinematic profiles separated injury states, we trained Random Forest (RF) classifiers using scikit-learn [9]. Before training, the KiMA pipeline applied pre-processing routines including outlier removal based on ±2 SD from the group median, individual gait-cycle segmentation, and concatenation of features across groups into a unified feature matrix. Each RF model comprised 100 decision trees using the Gini impurity criterion, with tree depth and maximum leaf nodes left unconstrained. Separate RF classifiers were trained for each pairwise comparison (Uninjured vs. 1 WPI, Uninjured vs. 3 WPI). Data were partitioned by randomized stratified sampling into a 75% training set and 25% held-out test set; accuracy and confusion matrices were computed on the test set only. Feature importances were ranked by Mean Decrease in Impurity (Gini importance) to identify the kinematic features contributing most to state separation.

### Animals husbandry

Adult C57BL/6J mice (2–4 months old, both sexes) were housed under standard conditions: 12 h light/dark cycle, ambient temperature 23–25 °C, and humidity 40–60%, with food and water available *ad libitum*. All animal protocols and experimental procedures were performed in accordance with institutional guidelines and were formally approved by the local Institutional Animal Ethics Committee.

### Thoracic spinal cord injury

A total of 30 mice were used for behavioral and kinematic evaluation. Beginning four days before surgery (Days −4 to −2), animals were habituated daily to all three behavioral apparatuses (Ladder/Rung, Foot Fall, CatWalk) to reduce handling-induced anxiety and establish baseline performance (Fig. 2a, 3a, 4a).

**Figure. 1:**
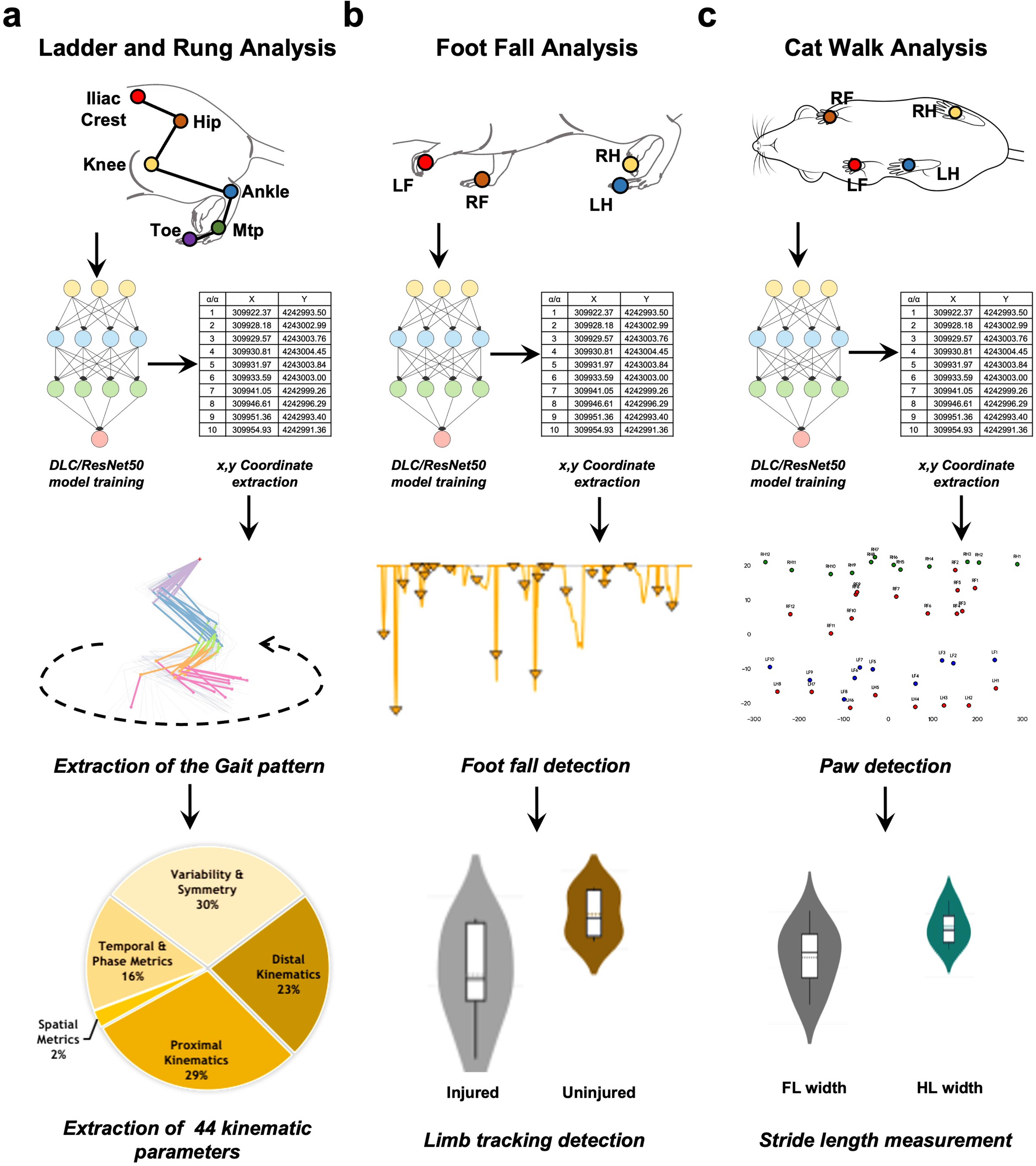
Modular workflow of the KiMA toolbox for automated mouse locomotor and kinematic profiling. (a) Ladder and Rung analysis pipeline. Video frames of mice traversing the apparatus are processed using DeepLabCut (DLC; ResNet-50 network architecture) to extract (x, y) coordinates from six anatomical landmarks along the hindlimb: iliac crest, hip, knee, ankle, metatarsophalangeal (MTP) joint, and toe. KiMA processes coordinate traces to extract full gait cycle patterns and quantify 44 distinct kinematic parameters divided across five functional classes: Distal Kinematics (23%), Proximal Kinematics (29%), Temporal & Phase Metrics (16%), Spatial Metrics (2%), and Variability & Symmetry metrics (30%). (b) Foot Fall analysis pipeline. Markerless labeling of four limb endpoints (LF: left forelimb, RF: right forelimb, LH: left hindlimb, RH: right hindlimb) followed by (x, y) coordinate extraction. KiMA detects footfall events and generates limb tracking profiles, enabling statistical comparison (e.g., violin plots) between injured (thoracic SCI) and uninjured states. (c) Cat Walk analysis pipeline. Ventral-view recordings are analyzed following DLC coordinate extraction for all four paws. KiMA performs automated paw detection to measure spatial stride parameters, including forelimb (FL) and hindlimb (HL) width metrics, to evaluate locomotor recovery post-treatment.

**Figure. 2:**
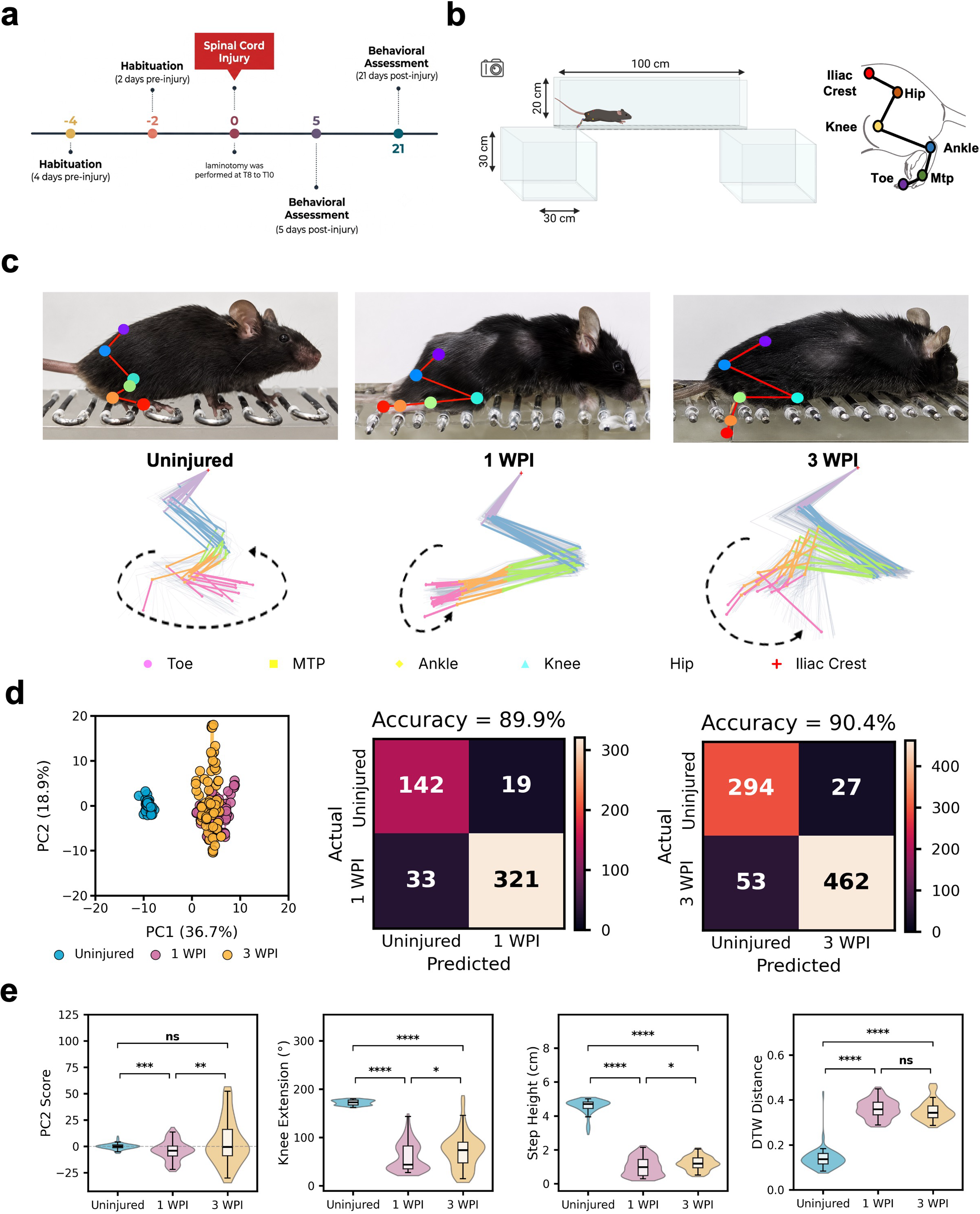
Quantitative kinematic profiling and machine learning classification of locomotor recovery post-spinal cord injury. (a) Experimental timeline. Schematic showing baseline habituation (Days -4 and -2), thoracic (T8–T10) spinal cord injury (Day 0), and post-injury behavioral assessment timepoints at 5 DPI (∼1 WPI) and 21 DPI (3 WPI). (b) Behavioral apparatus and joint labeling schematic. Dimensions of the overground walkway (100 cm length, 20 cm height, 30 cm elevation) with camera positioning and anatomical location of the six labeled hindlimb landmarks: iliac crest, hip, knee, ankle, metatarsophalangeal (MTP) joint, and toe. (c) Representative DLC tracking and gait stick plots. Upper panels display DLC markerless tracking on representative video frames from Uninjured, 1 WPI, and 3 WPI mice. Lower panels display corresponding stick-diagram reconstructions of hindlimb trajectories over full gait cycles, showing post-injury trajectory collapse and partial restoration at 3 WPI. (d) Multivariate profiling and machine learning classification. Left: Principal Component Analysis (PCA) plot of n = 90 datasets across 44 kinematic parameters showing distinct clustering of Uninjured (blue), 1 WPI (magenta), and 3 WPI (yellow) states along PC1 (36.7% variance) and PC2 (18.9% variance). Right: Confusion matrices and classification accuracy of Random Forest models predicting Uninjured vs. 1 WPI (89.9% accuracy) and Uninjured vs. 3 WPI (90.4% accuracy).(e) Quantification of primary kinematic features and gait dissimilarity. Violin/box plots comparing PC2 score, Knee Extension (°), Step Height (cm), and Dynamic Time Warping (DTW) distance across Uninjured, 1 WPI, and 3 WPI groups. Pairwise comparisons (Uninjured vs. 1 WPI) were evaluated using Student’s t-test; multi-group comparisons across all three stages were analyzed using one-way ANOVA with Tukey’s post-hoc test. Significance levels: ns = not significant (P > 0.05), *P < 0.05,P < 0.01, ***P < 0.001, ****P < 0.0001.

**Figure. 3:**
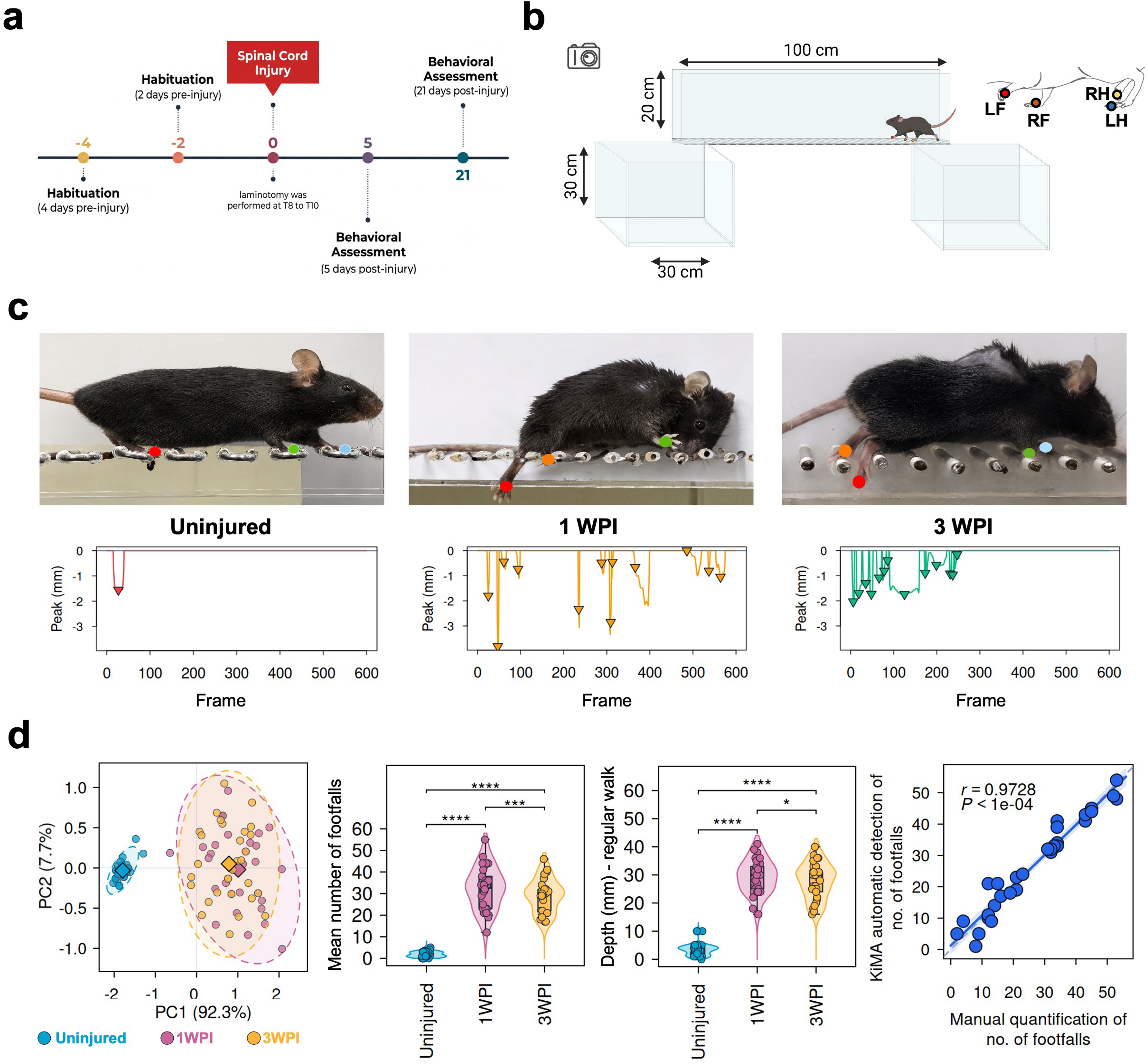
Automated detection and quantitative profiling of skilled paw placement deficits using the KiMA Foot Fall module. (a) Experimental timeline. Experimental schedule indicating habituation (Days -4 and -2), thoracic (T8–T10) spinal cord injury (Day 0), and post-injury behavioral assessments at 5 DPI (1 WPI) and 21 DPI (3 WPI). (b) Apparatus layout and limb labeling. Schematic of the elevated ladder rung apparatus (100 cm runway length, 20 cm height, 30 cm elevation) and anatomical labeling positions for the four limb extremities: left forelimb (LF), right forelimb (RF), left hindlimb (LH), and right hindlimb (RH). (c) Representative video tracking and footfall signal traces. Upper panels show markerless DLC limb tracking on representative frames from Uninjured, 1 WPI, and 3 WPI mice traversing the ladder. Lower panels show corresponding KiMA peak detection traces displaying negative vertical coordinate displacement (Peak in mm) representing footfall errors across video frames.(d) Multivariate grouping, footfall parameter quantification, and automated algorithm validation. From left to right: (1) PCA plot displaying clustering of Uninjured (blue), 1 WPI (magenta), and 3 WPI (yellow) datasets along PC1 (92.3% variance) and PC2 (7.7% variance).(2) Violin/box plot of mean number of footfalls across groups. (3) Violin/box plot of footfall depth (in mm) across groups. Non-parametric repeated measures were analyzed using the Friedman test (*P < 0.05, ***P < 0.001, ****P < 0.0001). (4) Regression plot showing strong positive correlation between KiMA automated footfall detection and manual expert scoring (r = 0.9728, P < 0.0001).

**Figure. 4:**
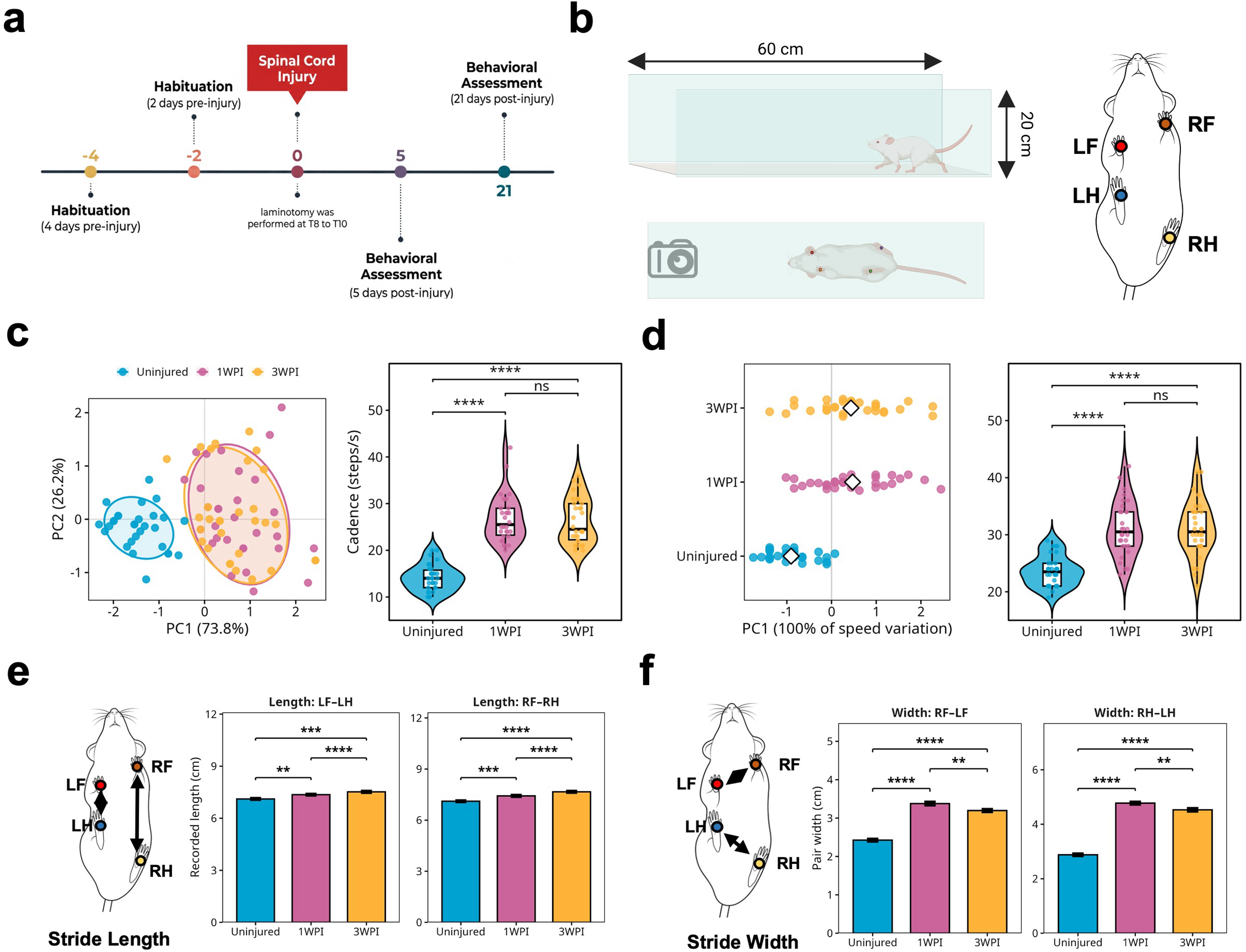
Ventral-view Cat Walk module analysis of cadence, speed, stride length, and stance width dynamics post-spinal cord injury. (a) Experimental schedule. Timeline outlining habituation (Days -4 and -2), thoracic (T8–T10) spinal cord injury (Day 0), and post-injury assessment days (5 DPI / 1 WPI and 21 DPI / 3 WPI). (b) Apparatus schematic and ventral paw tracking. Transparent walkway dimensions (60 cm length, 20 cm height) with bottom-mounted camera setup. Schematic highlights anatomical marker placements for markerless tracking of all four paws: left forelimb (LF), right forelimb (RF), left hindlimb (LH), and right hindlimb (RH).(c) Cadence profiling. Left: PCA plot displaying dataset clustering for Uninjured (blue), 1 WPI (magenta), and 3 WPI (yellow) across PC1 (73.8% variance) and PC2 (26.2% variance). Right: Box/violin plot of Cadence (steps/s) across timepoints. (d) Speed profiling. Left: PCA plot of speed variation (PC1 accounting for 100% of variance). Right: Box/violin plot of locomotion Speed (cm/s) across timepoints. (e) Stride Length quantification. Left: Schematic illustrating longitudinal stride length measurement between ipsilateral limb pairs (LF–LH and RF–RH). Right: Bar plots (mean ± SEM) displaying recorded stride lengths across Uninjured, 1 WPI, and 3 WPI groups. (f) Stride Width quantification. Left: Schematic illustrating lateral base-of-support (stride width) measurement across forelimb (RF–LF) and hindlimb (RH–LH) pairs. Right: Bar plots (mean ± SEM) displaying pair widths across timepoints. Multi-group comparisons across all three stages were analyzed using the non-parametric Friedman test; pairwise group comparisons were evaluated using Wilcoxon rank-sum tests with Holm adjustment for multiple testing. Significance levels: ns = not significant (P > 0.05), *P < 0.05,P < 0.01, ***P < 0.001, ****P < 0.0001.

On Day 0, mice were anesthetized with an intraperitoneal injection of ketamine (50 mg/ml) and xylazine (20 mg/ml) according to animal body weight. The animal is secured in a manual spinal stabiliser. A midline skin incision was made along the dorsal thorax region, and the paraspinal muscles were retracted to expose the vertebral column. A dorsal laminectomy was performed at T8–T10 to expose the spinal cord. A crush injury was then performed across the full width of the cord for 10 s using blunt end forceps [10], [11], [12]. Post-operatively, animals were kept on a heating pad until they recovered from anaesthesia, and meloxicam (analgesic) and enrofloxacin (antibiotic) were injected subcutaneously until 10 days post injury. Bladder expression was performed twice daily. Successful injury induction was verified by bilateral hindlimb paralysis. Behavioral and kinematic assessments were performed longitudinally from 5 to 21 days post-injury (DPI); primary analyses focused on 5 DPI (1 week post-injury, 1 WPI) and 21 DPI (3 WPI) to capture acute deficit and recovery, respectively.

### Behavioral apparatuses and video recording

Custom behavioural apparatuses were fabricated in-house using 3 mm transparent acrylic sheets to permit unobstructed video capture. The horizontal ladder measured 100 cm in length and 20 cm in height, elevated 30 cm above the bench surface (Fig. 2b). Side walls featured 2 mm-diameter holes drilled at 1 cm intervals to secure stainless steel rungs (2 mm diameter x 3 cm length), arranged at a regular 1 cm spacing. The CatWalk runway was build using 3 mm clear acrylic sheets, measuring 60 cm in length, 10 cm in width, and 20 cm in wall height (Fig. 4b).

All behavioral sessions were recorded at 1080p resolution and 30 frames per second (fps). For the Ladder & Rung and Foot Fall modules, the camera was mounted laterally, positioned 45 cm perpendicular to the apparatus to capture side-view hindlimb trajectories (Fig. 2b, 3b). For the CatWalk module, the camera was mounted beneath the transparent floor, facing upward to record paw placement, stride length, and stance width from a ventral perspective (Fig. 4b)).

### Blinded manual scoring of footfalls

To validate KiMA’s automated footfall detection, an independent manual scoring protocol was performed. two evaluators, blinded to experimental condition (uninjured, 1 WPI, 3 WPI), were trained to identify footfall errors, defined as partial or complete slippage of a limb extremity below the upper plane of the ladder rungs. Each evaluator was then given randomized ∼1-min video clips from the horizontal ladder task and asked to count all footfall events per trial and log the frame number of each error. Manually annotated frame indices were converted to frame-by-frame occurrence traces and compared against the peak-detection output of KiMA to compute the Pearson correlation between automated and manual counts (r = 0.9728, P < 0.0001; Fig. 3d).

### Statistics and reproducibility

Data are presented as mean ± SEM in summary plots and as violin/box plots showing medians, IQRs, and individual data points (n = 30 mice). Analyses were performed in Python using SciPy and scikit-learn. Normality was assessed with the Shapiro Wilk test. Multi-group comparisons across timepoints (Uninjured, 1 WPI, 3 WPI) were performed by repeated-measures one-way ANOVA with Tukey’s post-hoc test for parametric data, and by the Friedman test for non-parametric data (e.g., cadence, speed, footfall metrics). Pairwise comparisons between stages used Student’s t-test or Wilcoxon rank-sum tests with Holm adjustment for multiple testing. Agreement between automated KiMA detection and manual scoring was quantified with Pearson’s correlation coefficient (r). Multivariate structure was visualized with PCA on standardized (z-scored) features. Automated classification of injury and recovery states used Random Forest models (100 trees; Gini impurity) trained on a 75/25 stratified split, as described above.

## Results

### Automated end-to-end framework for mouse gait and footfall kinematic analysis using KiMA

We developed KiMA to enable high-throughput, automated quantification of motor recovery and gait deficits after thoracic spinal cord injury (SCI). KiMA combines deep-learning-based markerless tracking with a downstream analysis suite spanning three behavioral modules: Ladder/Rung, Foot Fall, and CatWalk (Fig. 1). For each module, locomotor videos are first processed with DeepLabCut (DLC) using a ResNet-50 backbone to extract frame-by-frame (x, y) coordinate matrices for each labeled landmark (Fig. 1a–c).

The Ladder & Rung module (Fig. 1a) targets six anatomical points along the hindlimb: iliac crest, hip, knee, ankle, metatarsophalangeal (MTP) joint, and toe. KiMA converts the raw coordinates into stick-plots, gait reconstructions and computes 44 kinematic parameters grouped into five categories: distal kinematics (10 parameters; MTP and ankle metrics), proximal kinematics (13 parameters; hip, knee, and pelvic tilt metrics), temporal and phase metrics (7 parameters; stance time, swing time, cadence), spatial metrics (1 parameter; stride length), and variability and symmetry (13 parameters; gait regularity, cadence consistency, stance/swing symmetry).

The Foot Fall module (Fig. 1b) tracks the four limb endpoints left forelimb (LF), right forelimb (RF), left hindlimb (LH), and right hindlimb (RH) for detection of skilled paw-placement errors. Users can isolate individual limbs to inspect spatial trajectories, automated footfall detection traces, and tracking quality. Comparing footfall counts between uninjured controls and thoracic SCI animals provides a sensitive readout of fine motor deficits and their recovery over time.

The CatWalk module (Fig. 1c) uses ventral-view video to record paw placement during overground locomotion. Tracking of the four paw markers supports automated stance detection and quantification of forelimb and hindlimb stride lengths, stance widths, and base of support. Together, the three modules provide complementary readouts of gross gait, skilled paw placement, and spatial footprint geometry within a single analysis framework.

### Tracking locomotor dynamics and recovery after spinal cord injury with KiMA

To test whether KiMA could resolve functional deficits and recovery after neurotrauma, we evaluated overground locomotion in a thoracic SCI mouse model (Fig. 2a). Animals were habituated on Days −4 and −2, underwent T8–T10 thoracic laminectomy and crush injury on Day 0, and were re-assessed at 1 WPI and 3 WPI (Fig. 2a). Locomotor trials were recorded on the 100 cm elevated ladder-free runway (Fig. 2b). Six hindlimb landmarks iliac crest, hip, knee, ankle, MTP joint, and toe were tracked frame-by-frame using DeepLabCut (ResNet-50 backbone) to produce continuous (x, y) coordinate traces (Fig. 2b, c). Stick-plot reconstructions revealed clear kinematic disruption at 1 WPI compared with uninjured baseline, including collapsed joint trajectories and loss of limb extension, with partial restoration of the cyclic gait geometry by 3 WPI (Fig. 2c).

To capture multivariate gait alterations, we performed Principal Component Analysis (PCA) across all 44 kinematic parameters (n = 90 trials; Fig. 2d). Uninjured animals formed a tight cluster, while 1 WPI and 3 WPI trials separated along PC1 (36.7% of variance) and PC2 (18.9% of variance), showing both acute divergence from baseline and a progressive shift toward the uninjured cluster during recovery (Fig. 2d). Using the same feature matrix, we trained Random Forest classifiers to test whether kinematic profiles alone could identify injury state. The classifiers achieved 89.9% accuracy in distinguishing Uninjured from 1 WPI (confusion matrix: 142 true Uninjured, 321 true 1 WPI) and 90.4% accuracy for Uninjured vs. 3 WPI (294 true Uninjured, 462 true 3 WPI) (Fig. 2d).

To identify the kinematic features driving this separation, we examined the top-loading variables on PC2 together with the Gini-ranked features from the classifier (Fig. 2e). PC2 scores dropped at 1 WPI (P < 0.001) and partially recovered by 3 WPI (P < 0.01). Knee extension showed the largest single change: extension range collapsed at 1 WPI (P < 0.0001 vs. uninjured) and partially recovered by 3 WPI (P < 0.05 vs. 1 WPI). Step height followed the same pattern (P < 0.0001 vs. uninjured; P < 0.05 for 3 WPI vs. 1 WPI). Dynamic Time Warping (DTW) distance, which measures overall deviation from an uninjured reference gait trajectory, rose sharply at 1 WPI (P < 0.0001) and remained elevated at 3 WPI (P < 0.0001 vs. uninjured), indicating that residual micro-gait irregularities persist even when the animal has regained overt locomotion (Fig. 2e).

To resolve the temporal structure underlying these changes, we plotted joint angles across the normalized gait cycle (0–100% of stride duration) for pelvis pitch, hip flexion, knee flexion, and ankle dorsiflexion (Fig. S1). Uninjured animals showed the expected phase-locked oscillations in all four joints. At 1 WPI, these oscillations were largely absent, with the range of motion flattened across all joints. At 3 WPI, joint recovery was heterogeneous: pelvis pitch and knee flexion remained flattened, while hip flexion and ankle dorsiflexion showed a baseline shift relative to 1 WPI, consistent with partial restoration of postural control and distal joint engagement without full recovery of proximal joint rhythm (Fig. S1).

### Tracking skilled paw placement and fine motor deficits after spinal cord injury with KiMA

Gross overground locomotion is largely supported by intraspinal locomotor circuits, whereas precise paw placement on challenging surfaces such as ladder rungs requires intact descending supraspinal input [13]. To test whether KiMA can quantify these fine motor deficits, we applied the Foot Fall module to horizontal ladder-rung walking after thoracic SCI (Fig. 3a).

Animals traversed the elevated ladder apparatus (100 cm long, elevated 30 cm) before injury (uninjured baseline), at 5 DPI (1 WPI), and at 21 DPI (3 WPI) (Fig. 3a, b). All four limb endpoints LF, RF, LH, and RH were tracked with DeepLabCut (ResNet-50 backbone) (Fig. 3b). KiMA then identified footfall events by peak detection on the vertical (y-axis) displacement of each limb, flagging drops of the paw below the rung plane as errors (Fig. 3c). Uninjured animals produced few footfall events, whereas 1 WPI animals showed frequent, deep downward deviations of the paws; deviation frequency and depth were both reduced by 3 WPI, though neither returned to baseline (Fig. 3c).

PCA on the extracted footfall parameters separated uninjured baseline from post-injury timepoints, with PC1 and PC2 capturing 92.3% and 7.7% of the variance, respectively (Fig. 3d). Non-parametric analysis across timepoints (Friedman test) confirmed strong effects of injury on both footfall count and depth (Fig. 3d). The mean number of footfalls per trial rose at 1 WPI compared with uninjured controls (P < 0.0001) and declined by 3 WPI (P < 0.001 vs. 1 WPI), remaining higher than baseline (P < 0.0001 vs. uninjured). Footfall depth (in pixels) followed the same trajectory: an increase at 1 WPI (P < 0.0001 vs. uninjured) followed by partial recovery at 3 WPI (P < 0.05 vs. 1 WPI; P < 0.0001 vs. uninjured) (Fig. 3d).

To validate the automated detection, we compared KiMA footfall counts against blinded manual scoring on the same clips. Automated and manual counts were tightly correlated (Pearson r = 0.9728, P < 0.0001; Fig. 3d), indicating that KiMA’s peak-detection routine reproduces expert scoring without the throughput cost.

### Ventral-view profiling of gait speed, stride geometry, and base of support with KiMA

To characterize spatial gait parameters and base of support after SCI, we applied KiMA’s CatWalk module (Fig. 4). Videos were recorded from below a transparent walkway (60 cm long, 20 cm high) to track the four paw endpoints LF, RF, LH, RH at baseline, 1 WPI (5 DPI), and 3 WPI (21 DPI) (Fig. 4a, b).

PCA on the temporal locomotion metrics separated uninjured animals from injured timepoints along PC1 (73.8% of variance) and PC2 (26.2% of variance) (Fig. 4c). Cadence (steps/s) at 1 WPI relative to uninjured controls (P < 0.0001, Wilcoxon rank-sum with Holm correction) and remained elevated at 3 WPI (P < 0.0001 vs. uninjured; n.s. vs. 1 WPI), consistent with a shift to shorter, faster steps rather than recovery of a normal step frequency (Fig. 4c). Walking speed followed the same pattern, with a sustained increase after injury (P < 0.0001 at both 1 WPI and 3 WPI vs. uninjured; Friedman test P < 0.0001) (Fig. 4d).

To quantify spatial footprint geometry, KiMA computed longitudinal (stride length) and lateral (stride width) inter-paw distances (Fig. 4e, f). Stride length between ipsilateral forelimb-hindlimb pairs (LF–LH and RF–RH) increased at 1 WPI (P < 0.01 for LF–LH, P < 0.001 for RF–RH vs. uninjured) and increased further at 3 WPI (P < 0.0001 for both pairs vs. uninjured) (Fig. 4e). Stride width the base of support between forelimb (LF–RF) and hindlimb (LH–RH) pairs widened at 1 WPI (P < 0.0001 vs. uninjured), and narrowed partially by 3 WPI (P < 0.01 vs. 1 WPI) while remaining wider than baseline (P < 0.0001 vs. uninjured) (Fig. 4f). Together, the persistent elevation of cadence, speed, and stride length, alongside partial recovery of stance width, indicates that animals adopt a compensatory gait strategy-shorter, faster, steps taken with a wider base of support, rather than returning to their pre-injury walking pattern by 3 WPI.

### Benchmarking KiMA against existing tools for automated footfall detection

To position KiMA’s performance relative to existing open-source tools, we benchmarked it against ALMA (Automated Limb Motion Analysis)[8], a recently published Python-based toolbox that operates on the same DeepLabCut pipeline and ResNet-50 backbone. Both tools were run on an identical set of 30 ladder-rung video samples spanning uninjured, 1 WPI, and 3 WPI animals, and their automated footfall counts were compared against blinded manual scoring by expert observers as the reference standard (Fig. 5a).

**Figure 5.**
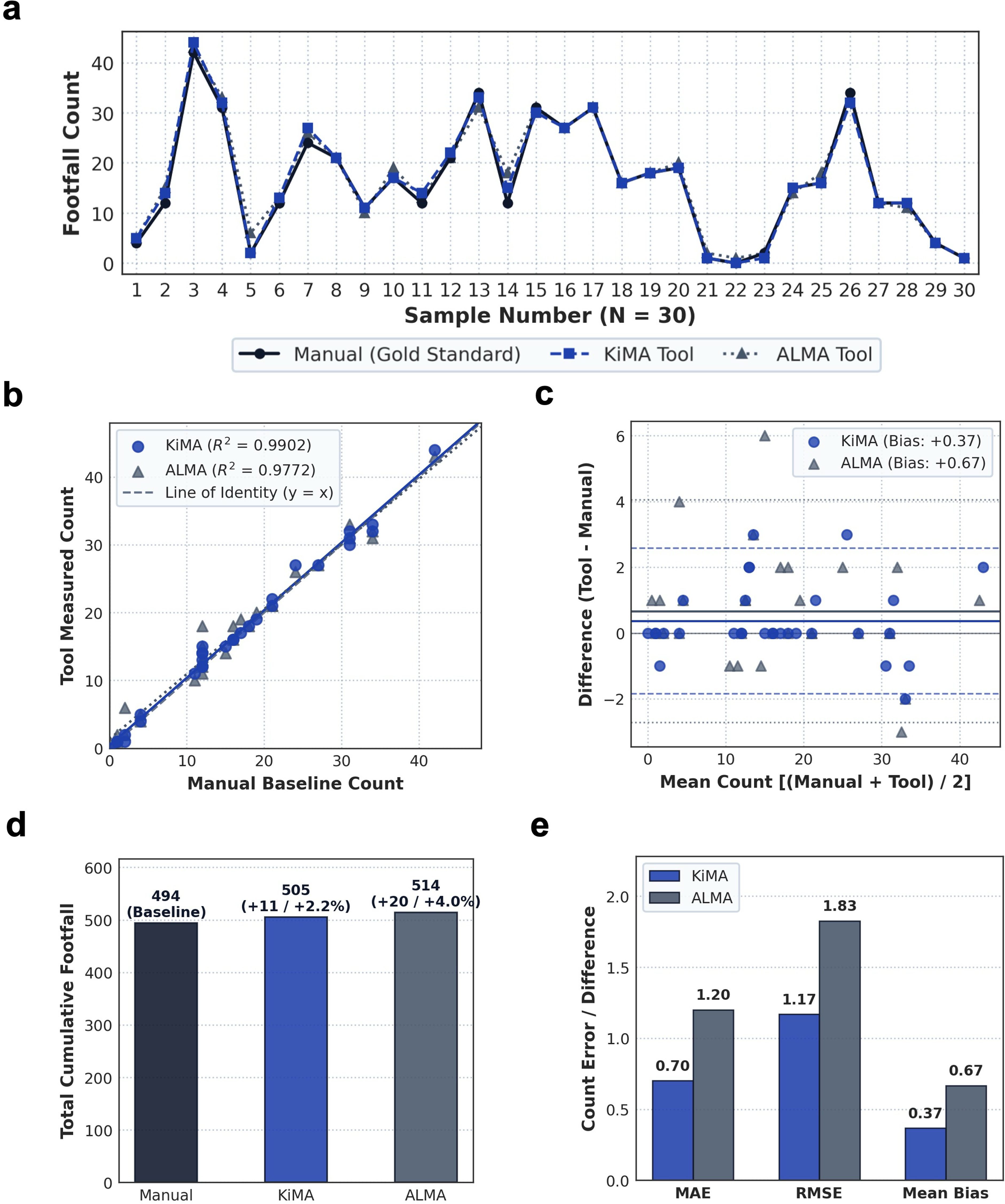
Comparison and evaluation of automated footfall counting tools against the manual standard baseline across 30 samples. (a) Sample-by-sample footfall count tracking across 30 distinct observation intervals comparing Manual counting (gold standard, solid line with circular markers), KiMA tool (dashed line with square markers), and ALMA tool (dotted line with triangular markers). (b) Accuracy scatter plot and linear regression analysis comparing tool-measured counts against manual baseline counts. The dashed diagonal line represents the line of identity (y = x). KiMA demonstrated stronger linear correlation (R2 = 0.9902) relative to ALMA (R2 = 0.9772). (c) Bland–Altman agreement plot displaying measurement differences (Tool count minus Manual count) against the mean of the two measurements for KiMA (blue circles) and ALMA (grey triangles). Solid horizontal lines indicate mean bias (+0.37 counts for KiMA; +0.67 counts for ALMA), while dashed lines denote the 95% Limits of Agreement (Mean ± 1.96 SD). (d) Cumulative total footfall count comparison across all 30 samples. KiMA recorded 505 footfalls (+2.23% bias; +11 counts) and ALMA recorded 514 footfalls (+4.05% bias; +20 counts) relative to the manual baseline of 494 counts. (e) Performance error metrics breakdown comparing Mean Absolute Error (MAE), Root Mean Squared Error (RMSE), and Mean Bias for KiMA and ALMA.

Across the 30 samples, KiMA’s per-sample footfall counts tracked the manual reference more closely than ALMA’s (Fig. 5a). Linear regression against the manual reference gave a slope near unity and a higher coefficient of determination for KiMA (R² = 0.9902) than for ALMA (R² = 0.9772), corresponding to a 2.33-fold reduction in residual variance (Fig. 5b).

Bland–Altman analysis of measurement agreement showed a systematic positive bias for both tools relative to manual counting, but the bias was smaller for KiMA (+0.37 counts) than for ALMA (+0.67 counts), a 45% reduction in mean bias (Fig. 5c). The 95% limits of agreement were also narrower for KiMA, indicating more consistent performance across samples.

Aggregated across all 30 samples, the manual reference yielded 494 footfalls in total. KiMA detected 505 footfalls (+2.23% bias, +11 counts), while ALMA detected 514 (+4.05% bias, +20 counts) — a 44.9% reduction in cumulative systematic error for KiMA relative to ALMA (Fig. 5d). Per-sample error metrics followed the same pattern: mean absolute error (MAE) was 42% lower for KiMA (0.70 vs. 1.20 counts) and root mean squared error (RMSE) was 36% lower (1.17 vs. 1.83 counts), indicating more accurate detection on each individual sample rather than only on the aggregate (Fig. 5e).

## Discussion

Objective, high-resolution quantification of locomotor function is a persistent limitation of preclinical spinal cord injury (SCI) research. The Basso Mouse Scale [1] and its rat predecessor, the Basso-Beattie-Bresnahan scale [1], remain the field’s default endpoints because they are simple and well-validated, but they were designed to capture gross recovery milestones and are known to be less sensitive at the extremes of the scale and to compress fine kinematic differences into a small number of ordinal steps [14], [15]. The horizontal ladder-rung task [14], [16] and grid walk [17] extend this coarse readout by exposing the animal to demands that depend on descending supraspinal input, chiefly the corticospinal tract[18], [19], but manual scoring of these tasks is time-consuming and observer-dependent [16]. Commercial gait platforms, principally CatWalk XT [3] and DigiGait [4], record a much larger number of spatiotemporal parameters and have been extensively validated against BMS and BBB in rodent SCI [2], [3], [20]. Their limitations are practical rather than technical, including high acquisition cost, proprietary analysis pipelines, and disagreement in the literature about which of the >100 CatWalk parameters best track SCI recovery [3], [20]. Markerless pose estimation with DeepLabCut [6] and SLEAP [7] has removed the tracking bottleneck, but converting per-frame coordinates into interpretable kinematic parameters is still non-trivial and has driven a second generation of downstream toolboxes, including GAITOR [21], ALMA [8], Visual Gait Lab [22], Anipose for 3D reconstruction [23], the MotoRater analysis pipeline [24], and unsupervised behavior segmenters such as B-SOiD [25], VAME [26], and Keypoint-MoSeq [27]. All of these are useful, and each addresses part of the problem, but with the exception of the commercial systems they are Python packages that require local installation, environment management, and, in most cases, a downstream step in GraphPad Prism or R for the statistics reviewers expect to see. KiMA was built to close this specific gap.

KiMA is a browser-based analysis suite that consumes standard DeepLabCut output and returns the parameters and statistics typically reported in rodent SCI studies. Its three modules (Ladder & Rung, Foot Fall, and CatWalk) are aligned with tasks that dissect distinct levels of the motor hierarchy. Overground gait is largely supported by lumbar central pattern generators and can persist even after severe supraspinal disconnection[28], [29], while ladder-rung placement is a canonical readout of descending cortico- and rubrospinal control [18], [19], [30]. In a thoracic crush model, the multivariate profile extracted by KiMA reproduced the expected pattern of acute deficit and partial recovery: PCA on 44 kinematic features separated uninjured, 1 WPI and 3 WPI animals along two principal axes, and Random Forest classifiers reached 89.9–90.4% accuracy. These figures are consistent with earlier deep-learning-based classifiers of rodent gait after neurotrauma (85–90% accuracy for stroke recovery from runway kinematics) [31] and with comparable ranges reported for clinical iSCI classification from motion-capture data [32], [33]. The knee-extension, step-height, and Dynamic Time Warping distance features that emerged as leading Gini-ranked contributors overlap with the joint-angle collapse and residual micro-gait irregularity reported in previous kinematic profiles of thoracic SCI [34]. The Foot Fall module, benchmarked against blinded manual scoring on the same clips, achieved a Pearson r of 0.973, in line with published inter-rater agreements for expert ladder scoring[14], [16]. Against ALMA, the most directly comparable open-source tool [16], KiMA showed modestly better agreement with the manual reference (R² 0.99 vs. 0.98; MAE 0.70 vs. 1.20; mean bias +0.37 vs. +0.67 counts on 30 videos). We interpret this gap as an implementation-level difference in the peak-detection routine rather than a fundamental architectural advantage, since both tools consume the same ResNet-50-based DLC coordinates. Both tools also over-count relative to human scorers, which suggests the residual disagreement lives at near-threshold events that a human observer resolves through context that a purely y-coordinate peak detector cannot access.

Several limitations of the present work merit explicit statement. First, KiMA is a 2D toolbox; joint angles are estimated from lateral- or ventral-view single-camera video and cannot capture out-of-plane rotation, which for hindlimb kinematics is a known source of error that 3D pipelines such as Anipose [23] and DANNCE [35] were built to address. For groups already investing in multi-camera setups, a 3D approach remains the more accurate option, and integrating a 3D-input pathway into KiMA is a natural next step. Second, KiMA’s outputs are only as good as the upstream DLC network. Jittery tracking or body-part swaps propagate directly into the kinematic parameters, and the same warning applies to every DLC-derived toolbox in this family [8], [23], [25]. Third, our benchmarking is scoped to footfall detection on a single lesion model and 30 clips; head-to-head comparison of joint-angle extraction, stride segmentation, and PCA/classifier behavior against ALMA [8], GAITOR [21], [22]Visual Gait Lab [22], and the CatWalk XT [36][3] standard remains open work, as does independent validation in other SCI models (contusion, hemisection) and in adjacent injuries (stroke, peripheral nerve). Fourth, our Random Forest classifier operates on trial- or cycle-level features and, in its current form, does not enforce animal-level separation between training and test folds, an issue every ML-based rodent behavior classifier has to confront explicitly [8][31]. Finally, our multi-variate approach follows the growing consensus that no single CatWalk or kinematic parameter faithfully captures SCI recovery, and that combining features via PCA or supervised learning improves sensitivity to spontaneous and treatment-driven improvement. Positioned in the current tool ecosystem, KiMA is not a replacement for CatWalk XT’s dedicated hardware, nor for full 3D reconstruction, nor for the unsupervised segmentation approaches that address a different question (what behavioral syllables exist?). It is intended as a low-friction analysis layer on top of DeepLabCut for laboratories that already run standard rodent behavioural tasks and want the kinematic quantification, cohort statistics, and multivariate visualisation in one place, in a browser, without a Python environment.

## Conclusions

We developed KiMA, a browser-based analysis suite that takes DeepLabCut coordinate files and returns the joint angles, spatiotemporal parameters, footfall events, and multivariate summaries used to quantify locomotor recovery in rodent SCI. In a thoracic crush model, KiMA resolved the expected pattern of acute deficit and partial recovery across three complementary tasks (Ladder & Rung, Foot Fall, and CatWalk), separated injury states via PCA, and reached 89–90% classification accuracy from kinematic features alone. Automated footfall detection agreed with blinded manual scoring at r = 0.973, and matched or modestly exceeded ALMA on the same dataset. KiMA is not a replacement for dedicated 3D reconstruction, commercial gait hardware, or unsupervised behavior segmentation; each of those addresses a different question. What it provides is an accessible analysis layer on top of DeepLabCut for the standard 2D behavioral tasks that most SCI laboratories already run, with the visualization and cohort statistics built in. The application, source code, and documentation are openly available at https://kimasuite.in and https://github.com/mano2991/KiMA, and we anticipate that lowering the setup cost of quantitative gait analysis will make kinematic readouts a more routine part of preclinical SCI studies.

## Ethics approval and consent to participate

All animal procedures were performed in accordance with the guidelines of and with prior approval from the Institutional Animal Ethics Committee (IAEC).

## Author Contributions

MK was responsible for conceptualization, investigation, methodology, data curation, software development, formal analysis, data interpretation, and writing of the original draft. NRS was responsible for data curation, software development, formal analysis. IV contributed to conceptualization, investigation, methodology, and writing review and editing.

## Data availability statement

The code supporting the findings of this study is publicly available at https://github.com/mano2991/KiMA and https://kimasuite.in/. The raw data are avilable at 10.6084/m9.figshare.33162509

## Acknowledgments

We thank CSIR CCMB for providing research facilities and institutional support. We also thank Dhanuush Balakannan, Athul Narayan P. S., Anisha S. Menon, Dr. Prakash Chermakani, and Yogesh Sahu for their technical assistance, Debugging the tool and support with the surgical procedures. We are grateful to the members of the Animal House Facility for providing the animals and assisting with animal care.

## Supplementary Figure

**Supplementary Figure. S1:**
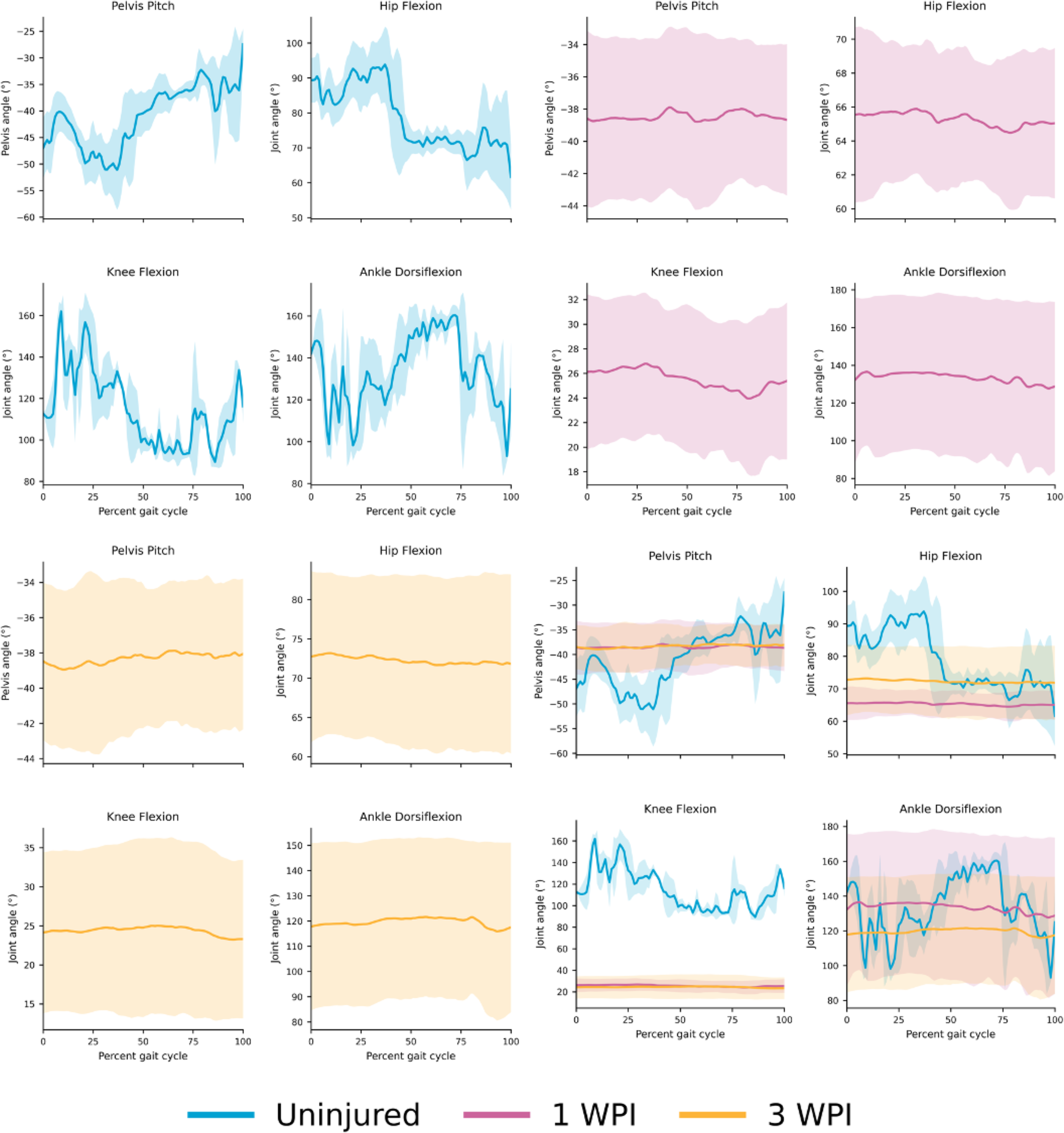
Continuous joint angle kinematic trajectories across normalized gait cycles post-spinal cord injury. Frame-by-frame joint angle profiles calculated from extracted (x, y) coordinates, normalized across full gait cycles (0% to 100% percent gait cycle). Panels display mean trajectory curves (solid lines) with shaded error bands representing standard deviation across Pelvis Pitch (°), Hip Flexion (°), Knee Flexion (°), and Ankle Dorsiflexion (°) for:Top-left panel (Blue): Uninjured baseline controls showing physiological dynamic angular range and cyclic oscillation. Top-right panel (Pink): 1 Week Post-Injury (1 WPI), illustrating severe trajectory flattening and loss of joint range of motion. Bottom-left panel (Yellow): 3 Weeks Post-Injury (3 WPI), showing persistent loss of cyclic amplitude. Bottom-right panel (Merged overlay): Direct comparative overlay of Uninjured (blue), 1 WPI (pink), and 3 WPI (yellow) trajectories. Merged traces highlight significant dynamic kinetic collapse post-SCI, with observable baseline shifts in Hip Flexion and Ankle Dorsiflexion between 1 WPI and 3 WPI despite unrecovered Pelvis Pitch and Knee Flexion profiles.

