## Supplementary figures and images for "KiMA: Kinematic Motion Analysis for Spinal Cord Injury Research"

### Supplementary Figure S1

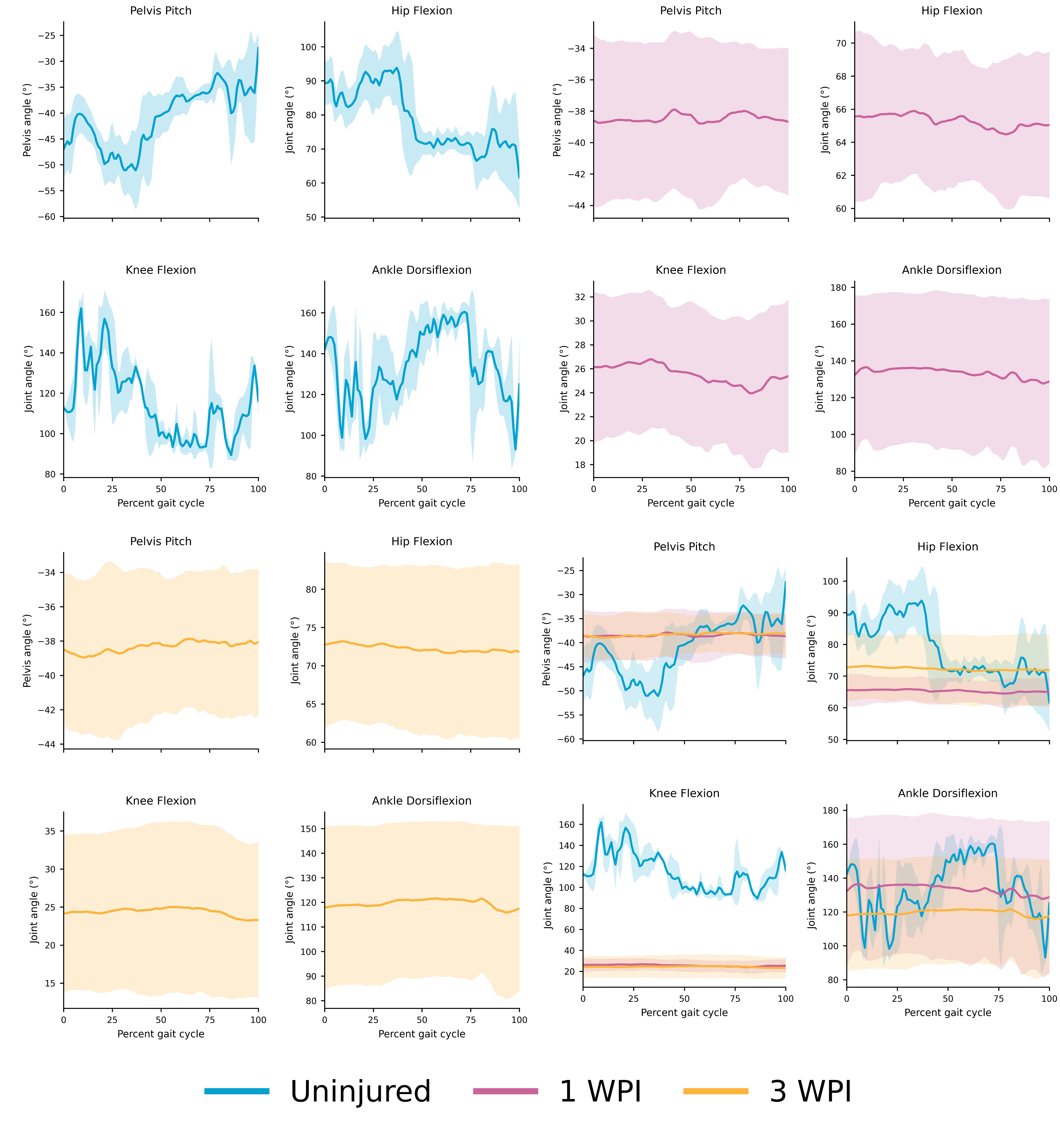
